# Posterior parietal cortex represents sensory history and mediates its effects on behavior

**DOI:** 10.1101/182246

**Authors:** Athena Akrami, Charles D. Kopec, Mathew E. Diamond, Carlos Brody

## Abstract

Many models of cognition and of neural computations posit the use and estimation of prior stimulus statistics^1–4^: it has long been known that working memory and perception are strongly impacted by previous sensory experience, even when that sensory history is irrelevant for the current task at hand. Nevertheless, the neural mechanisms and brain regions necessary for computing and using such priors are unknown. Here we report that the posterior parietal cortex (PPC) is a critical locus for the representation and use of prior stimulus information. We trained rats in an auditory Parametric Working Memory (PWM) task, and found that rats displayed substantial and readily quantifiable behavioral effects of sensory stimulus history, similar to those observed in humans^5,6^ and monkeys^7^. Earlier proposals that PPC supports working memory^8,9^ predict that optogenetic silencing of this region would impair behavior in our working memory task. Contrary to this prediction, silencing PPC significantly improved performance. Quantitative analyses of behavior revealed that this improvement was due to the selective reduction of the effects of prior sensory stimuli. Electrophysiological recordings showed that PPC neurons carried far more information about sensory stimuli of previous trials than about stimuli of the current trial. Furthermore, the more information about previous trial sensory history in the neural firing rates of a given rat’s PPC, the greater the behavioral effect of sensory history in that rat, suggesting a tight link between behavior and PPC representations of stimulus history. Our results indicate that the PPC is a central component in the processing of sensory stimulus history, and open a window for neurobiological investigation of long-standing questions regarding how perception and working memory are affected by prior sensory information.

---

Finding long-term regularities in the environment, and exploiting them, is a critical brain function in a complex yet structured world. But little is known about the neural mechanisms involved in estimating these regularities or their impact on memory. The history of sensory stimuli affects working memory (WM)^10,11^ and many other tasks involving sensory percepts^12,13^. One salient example, discovered over a century ago^14^ and repeatedly observed in human cognition^5,14,15^ is “contraction bias,” in which the representation of a stimulus held in working memory shifts towards the center of the distribution of stimuli observed in the past (the “prior distribution”). Despite the ubiquity of this phenomenon, and much psychophysical and theoretical research into the use and effects of prior stimulus distributions^2,3^, the neural mechanisms of contraction bias have not been identified.

Based on previous work using somatosensory stimuli^6^, and inspired by parametric working memory (PWM) tasks in primates^7^, we developed a computerized protocol to train rats, in high-throughput facilities, to perform a novel auditory PWM task (behavioral shaping code at <u>http://brodylab.org/auditory-pwm-task-code)</u>. PWM tasks involve the sequential comparison of two graded (i.e., analog) stimuli separated by a delay of a few seconds, here auditory pink noise stimuli, ‘s_a_’ and ‘s_b_’; rats were rewarded for correctly reporting which of the two was louder (**Fig. 1a**). Following ref.^16^, the set of [s_a_, s_b_] pairs used across trials in a session was chosen so that neither stimulus alone contained sufficient information to solve the task (**Fig. 1b**). As with any magnitude discrimination task, the smaller the difference between stimuli, the harder the task (**Fig. 1c**).

**Figure 1.**
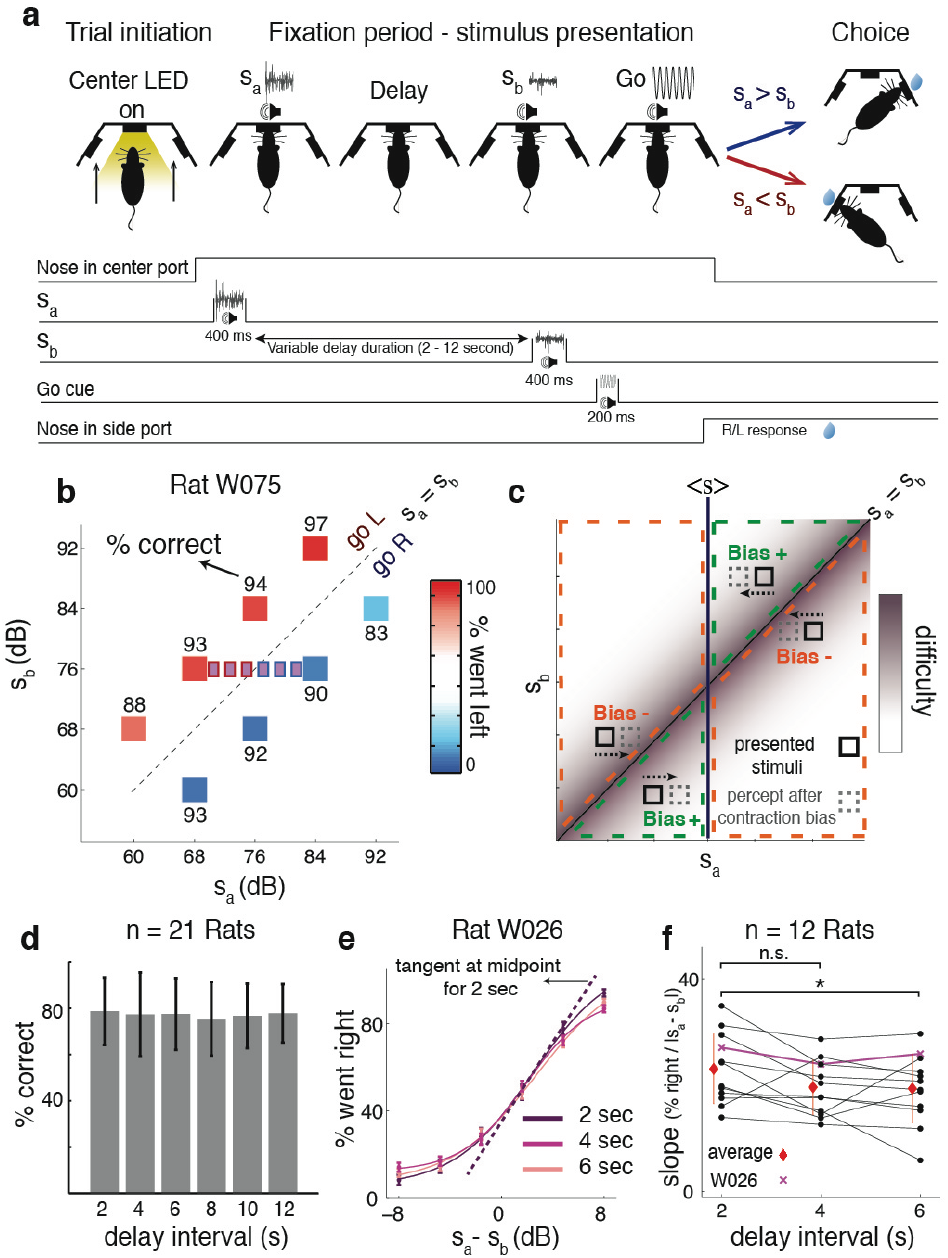
Rat performance and contraction bias. **a**, Rats compared two sequentially presented auditory stimuli ‘s_a_’ and ‘s_b_’, separated by a delay, and were rewarded for correctly reporting, through their choice of left or right nosepoke, which one was louder. **b**, Set of [s_a_, s_b_] pairs used in a session. On each trial, one randomly selected pair was presented. Small purple squares are stimuli used in a subset of sessions to assess performance at psychometric threshold. **c**, Contraction bias proposes that presented stimuli (black boxes) drive behavior as if s_a_ were closer (dashed boxes) to the average stimulus <s> (vertical midline) than its actual value. For some [s_a_, s_b_] pairs, this decreases their difference, and thus impairs performance (Bias-, red), while for others it has the opposite effect (Bias+, green). **d**, Overall average performance as a function of delay duration (n=21 rats, mean +/− STD over subjects). **e**, Psychometric curves for one example rat (n=120 sessions, mean +/− SEM over sessions, fits to a four-parameter logistic function; Methods). **f**, Midpoint tangent line slopes for the psychometric curves for each animal. These are significantly greater (reflecting better performance) at delays of 2 versus 6 seconds (two sample t-test: 2 vs. 4 sec: p = 0.051; 2 vs. 6 sec: p = 0.045; 4 vs. 6 sec: p = 0.86). Red, average and STD over rats.

Classical contraction bias^5^ argues that during the delay, the memory of the magnitude of s_a_ drifts towards the mean of all stimuli presented (**Fig. 1c**, vertical line “<s>“). Consequently, on those pairs in which s_a_ drifts away from the high difficulty s_a_=s_b_ diagonal, the memory of s_a_ becomes more distinct from s_b_, so contraction bias would improve performance (“Bias+”, **Fig. 1c**). In pairs where s_a_ drifts towards the diagonal, performance would decrease (“Bias−”). This predicted pattern can be seen in our rat behavior (**Fig. 1b**, high % correct for Bias+ stimulus pairs [s_a_=84, s_b_=92] and [s_a_=68, s_b_=60], lower % correct for Bias-stimulus pairs [s_a_=60, s_b_=68] and [s_a_=92, s_b_=84]). The same pattern has been observed in monkeys^7^ and humans (**Extended Data Fig 1d-e**, and^5,6,17^). History-dependent effects are likely adaptive in the natural world, where there are many long-term regularities. But in our laboratory task, in which each trial is generated independently, such biases, on average, produce suboptimal performance. The rats’ overall performance was robust and similar across delay intervals from 2-12 s (**Fig. 1d**; see **Extended Data Fig.1 b,c** for performance over learning). In some sessions, we used stimulus pairs that were closely spaced along s_a_ (**Fig. 1b**) to measure the psychometric discrimination threshold. This worsened slightly, but significantly, at longer delay periods (**Fig. 1e,f**).

In contrast to the small effects on overall performance from varying the delay interval (**Fig. 1d**), sensory history had a strong effect on performance. As quantified below, this effect was stronger than the well-documented influence of previous rewards and choices ^18–20^ (see also **Extended Data Fig.1g-i**). In trials that followed a rewarded right-choice trial, and thus holding previous choice and reward fixed, **Fig. 2a, left** shows that the smaller the previous trial’s stimuli, the greater the percentage of leftwards choices in the current trial (slope = −3.06 percent per decibel, linear fits to “% left - average”, p < 0.0001; see **Extended Data Fig. 5a** for slopes from n=1,..,7 trials back). This is consistent with a contraction bias in which the estimate of <s> is weighted towards recent stimuli^17^, because small recent values make the current s_a_ more likely to be perceived as small, increasing the likelihood of an “s_a_ < s_b_ (left)” response. **Fig. 2a, right** shows the same effect occurred across all combinations of current and previous trial stimuli from our standard stimulus set (|s_a_‒s_b_| fixed at 8 dB; see **Extended Data Fig. 3** for n=1,..,5 trials back, and **Extended Data Fig. 4** for controlling for action and reward). Similar effects were found in human auditory (**Fig. 2b**) or tactile (**Fig. 2c**) versions of the task, and increased for larger delay intervals^21,22^ (**Extended Data Fig. 2**). To simultaneously take into account effects across multiple previous trials of the history of rewards, choices, and stimuli, we fit logistic regression models with these variables as regressors, and compared the performance of a variety of such models on cross-validation data (**Fig. 2d**; Methods; **Extended Data Fig. 6**). Consistent with human data^17^, short-term (last two trials) sensory history had strong effects on behavior. In addition, our large dataset revealed a smaller but nevertheless important effect of longer-term (average of last few tens of trials) sensory history (**Fig. 2e**, **Extended Data Fig. 5b**). It has been suggested^21,23–27^ that sensory history does not add a behavioral bias independent of working memory, but instead produces a value of s_a_ in working memory that is a weighted average of the current stimulus and sensory history. In weighted averages, weights sum to one. Consistent with that suggestion, we found that constraining the sum of regression weights on the current trial’s s_a_ stimulus plus weights on previous sensory stimuli to equal one, thus removing one free parameter from the model, produced the best performance on cross-validation data (**Fig. 2e**, red; see **Extended Data Fig. 6** for all models and comparisons; the best-fitting model, for every individual animal, had the regressors in **Fig. 2d**). Sensory history was essential in accounting for behavior (**Fig. 2f,g**): examining the weights in the regression model (**Fig. 2h**) shows that those for sensory history are significantly larger than those for correct side history (p<0.001), and have a bigger impact on behavior (**Extended Data Fig. 6g**).

**Figure 2.**
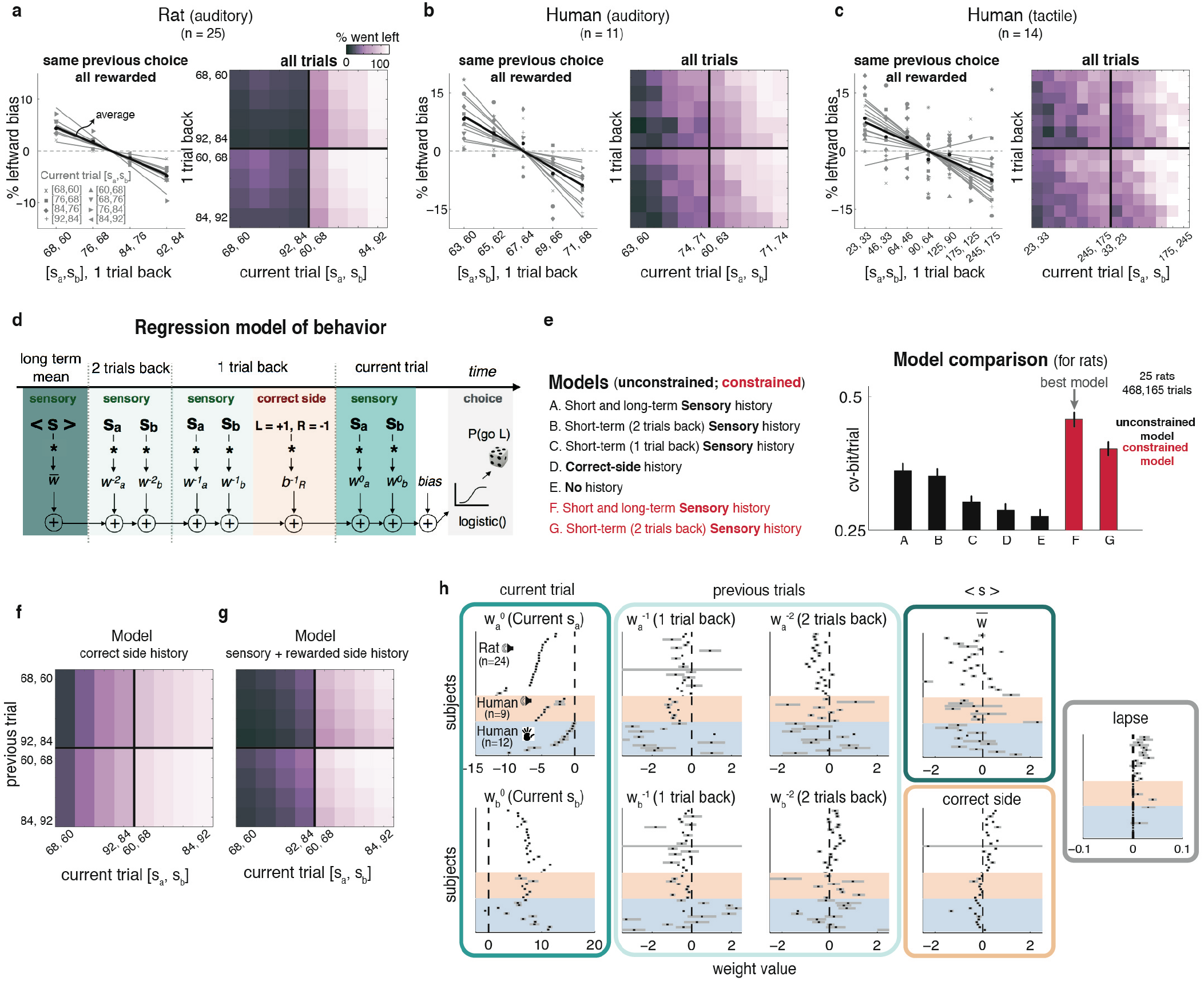
Sensory history biases behavior. **a**, Left: % trials went left - average went left, as a function of previous trial’s stimuli, for fixed previous trial response choice and reward. Right: % went left for each combination of current and previous stimuli; vertical modulation indicates previous trial effect. **b**, Format from (a), human subjects in auditory PWM task. **c**, Format from (a), human tactile data. **d**, Linear weighted sum of 9 regressors is used to predict the probability ratio log(*P_go Left_* / *P_go right_*); weights are fit to best-match training data, and are evaluated on left-out cross-validation data (Methods). Regressors: averaged stimuli over the last 20-50 trials (excluding last two); stimuli from last two trials; correct side on last trial (i.e., side baited with reward; when *b*^−1^_*R*_ is positive, this increases probability of going towards previously baited side, i.e., win-stay/lose-shift); current trial stimuli; overall side bias. **e**, Evaluation of model variants with different regressors (Methods; **Extended Data Fig. 6**). Best model has regressors shown in (d). Leftmost (A): Regressors as in (d). Moving rightwards, regressors progressively removed: long-term sensory history (B); last two trials’ stimuli (short-term history, C and D); previous trial correct side (E). Next two models ( red) have the same regressors as first two, but weights on current trial’s sa plus all previous sensory stimuli weights are constrained to add to 1, removing one free parameter. **f**, Poor match to data in (a) from predictions of a model with current trial and previous trial’s correct side regressors only. **g**, As in (f), now including sensory history regressors, greatly improves match to data. **h**, Black ticks, best-fit parameter values, per subject; gray bars, 95% CIs. All panels sorted based on value of W^0^_a_.

The PPC has been proposed as critical for working memory (^8,9,28^ but see ^29,30^),so we examined its role in our task. We injected bilaterally an AAV virus that drives expression of the light-activated inhibitory opsin halorhodopsin eNpHR3.0, under the CaMKIIa promoter (center of injection, AP -3.8 mm and ML 2.5 mm from Bregma, **Fig 3a, Extended Data Fig. 7a**). Optical fibers were inserted at the centers of injection sites to deliver laser illumination, and we inactivated PPC during a randomly chosen 20% of trials. To best probe for any small effects, we included psychometric stimuli (**Figs. 1b** and **3b**).

**Figure 3.**
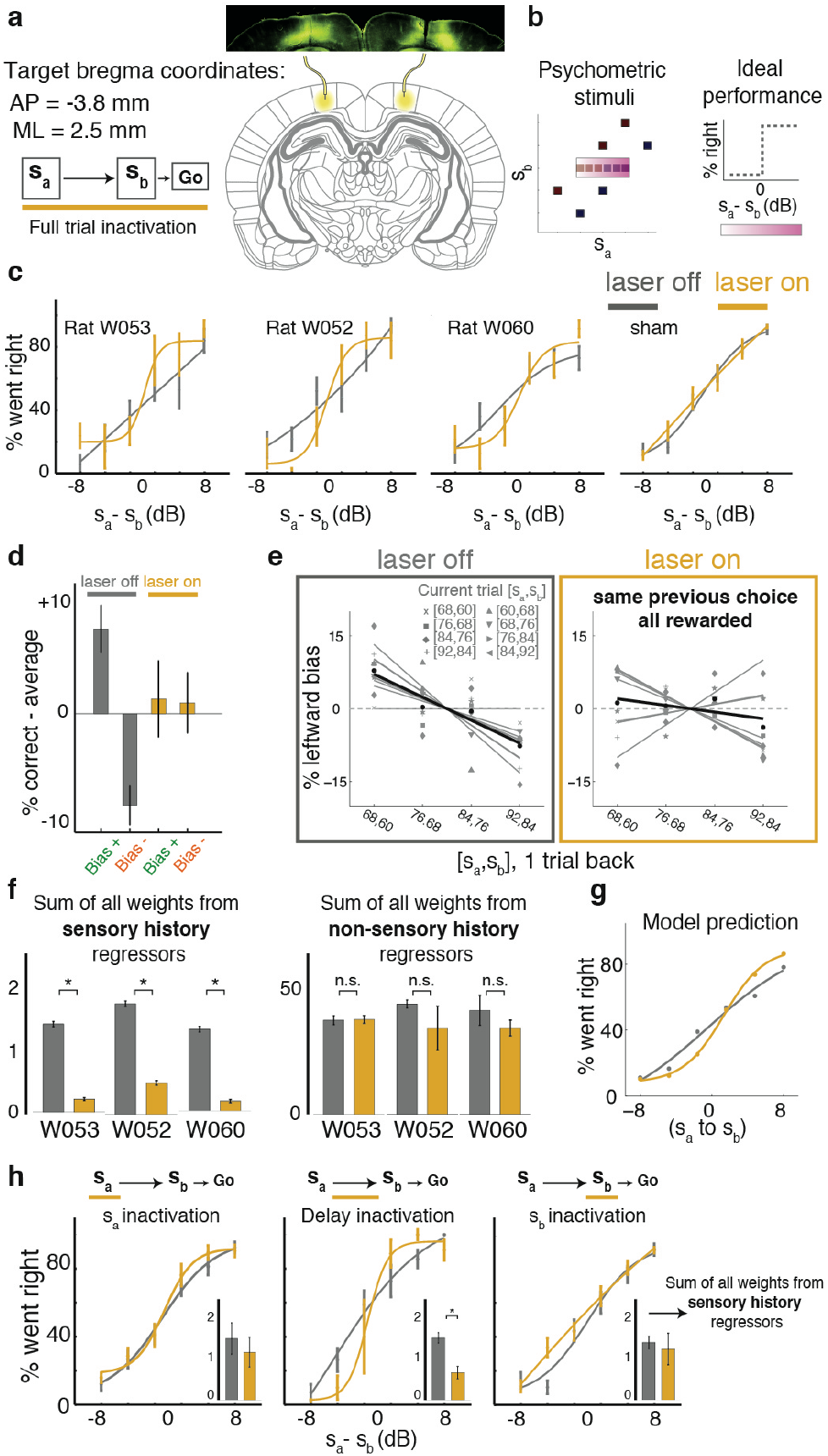
PPC is specifically necessary for behavioral effect of previous sensory stimuli. **a**, Schematic of virus injection and full trial inactivations. Brain image is taken from^31^ **b**. Stimuli included our standard [s_a_,s_b_] set (black) plus psychometric pairs (purple). Right: ideal psychometric performance. **c**, Psychometric curves for all animals were closer to ideal during PPC inactivation (yellow) trials than during control (grey) trials. Right, sham inactivation in rats with optic fibers but no virus had no effect (n=2). Error bars show SEM. d, %correct averaged across all Bias+ or all Bias-trials (**Fig. 1c**), relative to overall average performance. PPC inactivation eliminates the difference between Bias+ versus Bias− trials, or versus overall average (t-test, “Bias+” – “Bias −” significantly different from zero, laser off, p < 0.00001; laser on, p = 0.706; laser off vs on, p < 0.01; two-way ANOVA, interaction of laser on/off and “Bias+”/”Bias−”: p < 0.01). Error bars show SEM. **e**, The bias induced by previous stimuli is reduced under PPC inactivation (laser off: slope of −4.74, p = 0.0017; laser on: slope of −1.36, p = 0.42; laser on vs. off: p=0.044). **f**, PPC inactivation selectively reduces sensory history weights (left) in regression model of Fig. 2d. Error bars show 95% CI (n = 600, 200 iterations of 3fold CV; ‘*’ indicates p < 0.01, one-sided t-test). Right: sum of all other weights. See **Extended Data Fig. 9** for individual weights. **g**, Reducing sensory history weights in the model is sufficient to improve psychometric performance, comparable to experimental data (c). **h**, Similar to c, for inactivation during s_a_, or delay, or s_b_. Insets show sum of all sensory history regressors, as in f (n=600). Only inactivation of PPC during delay produces a significant effect (permutation test, s_a_, “laser off - on”: p = 0.17; delay : p < 0.0001; s_b_, : p = 0.18; s_a_ vs. delay: p = 0.03, s_b_ vs. delay: p = 0.02).

Expecting a performance impairment^8^, we were surprised to instead observe an improvement in psychometric performance in all animals tested (**Fig. 3c**). However, the effect was not simply an overall performance improvement: looking beyond the psychometric stimuli, while performance was indeed improved with respect to control on Bias- trials, PPC silencing instead impaired performance on Bias+ trials (**Fig. 3d**). Moreover, the difference between Bias+ and Bias- trials was eliminated, as was their difference from control average performance (**Fig. 3d; Extended Data Fig. 9a**). Similarly, bias as a function of the previous trial’s stimuli was markedly reduced (**Fig 3e**, laser off: p=0.42; laser on: p=0.0017; laser on vs. off: p=0.044, see **Extended Data Fig. 8** for impact on history matrices). Fitting our regression model separately to the set of laser on versus off trials, we found that PPC inactivation significantly reduced sensory history regression weights (**Fig. 3f**), and no other regression terms were significantly affected (individual weights in **Extended Data Fig. 9**). A model with reduced sensory history effects as in Fig. 3f was sufficient to reproduce the experimental data (**Fig. 3g**).

Thus PPC silencing appeared to have no impact on working memory but specifically and substantially reduced sensory history effects. This reduction did not persist into future trials (**Extended Data Fig. 8**), and occurred with PPC inhibition during the WM delay, but not during s_a_ or s_b_ (**Fig. 3h**), suggesting a focused role for the PPC in the interaction between previous stimuli and the current trial’s working memory.

To examine whether signatures of sensory history are present in the region whose inactivation appears to cancel history effects, we recorded extracellularly during task performance, from 936 units in five animals implanted with microwire arrays, targeting PPC. Neurons with a mean firing rate below 2 Hz were discarded (Methods), so 361 units were analyzed. Most cells were similar to the example cell in **Fig. 4a**; their firing rates during the working-memory delay did not distinguish between values of s_a_ held in memory, and therefore did not carry information about it. Instead, robust information about the sensory stimulus pair appeared approximately 1 second after the trial had terminated, during the inter-trial-interval (ITI). We used mutual information (MI; Methods) to quantify the amount of information, in neuronal firing rates, about which [s_a_ s_b_] sensory stimulus pair was presented (**Fig. 4b,c**; see **Extended Data Fig. 10** for MI about other components, including current and previous choices, rewards and s_a_ alone). During the ITI before a new trial, a large fraction of PPC neurons carried significant information about the previous trial stimuli (22% of analyzed neurons; **Fig. 4c**). A smaller fraction of cells continued to code the previous trial’s stimuli into the new trial’s start, and throughout the new trial’s WM delay (**Fig. 4c, Extended Data Fig. 10b**). We computed the fraction of neurons with significant MI about the previous trial’s stimuli, during the ITI (**Fig 4d**) or the current trial (**Fig 4e**), and compared this to the strength of the rat’s sensory history behavioral bias (**Fig 4f**). During the new trial, but not during the ITI, these two measures were perfectly correlated (**Fig 4g**, Spearman’s rank correlation r=1, p<0.01 during current trial; r=0.3, p=0.68 during ITI; p<0.00001 ITI vs. full current trial from Steiger’s Z-test). This suggests first, a tight link between sensory history representations in PPC and history biases, and second, that the PPC history representation is used during or shortly after the presentation of the new s_a_ (**Fig. 4h**), consistent with the idea that contraction bias affects the working memory representation of s_a_.

**Figure. 4.**
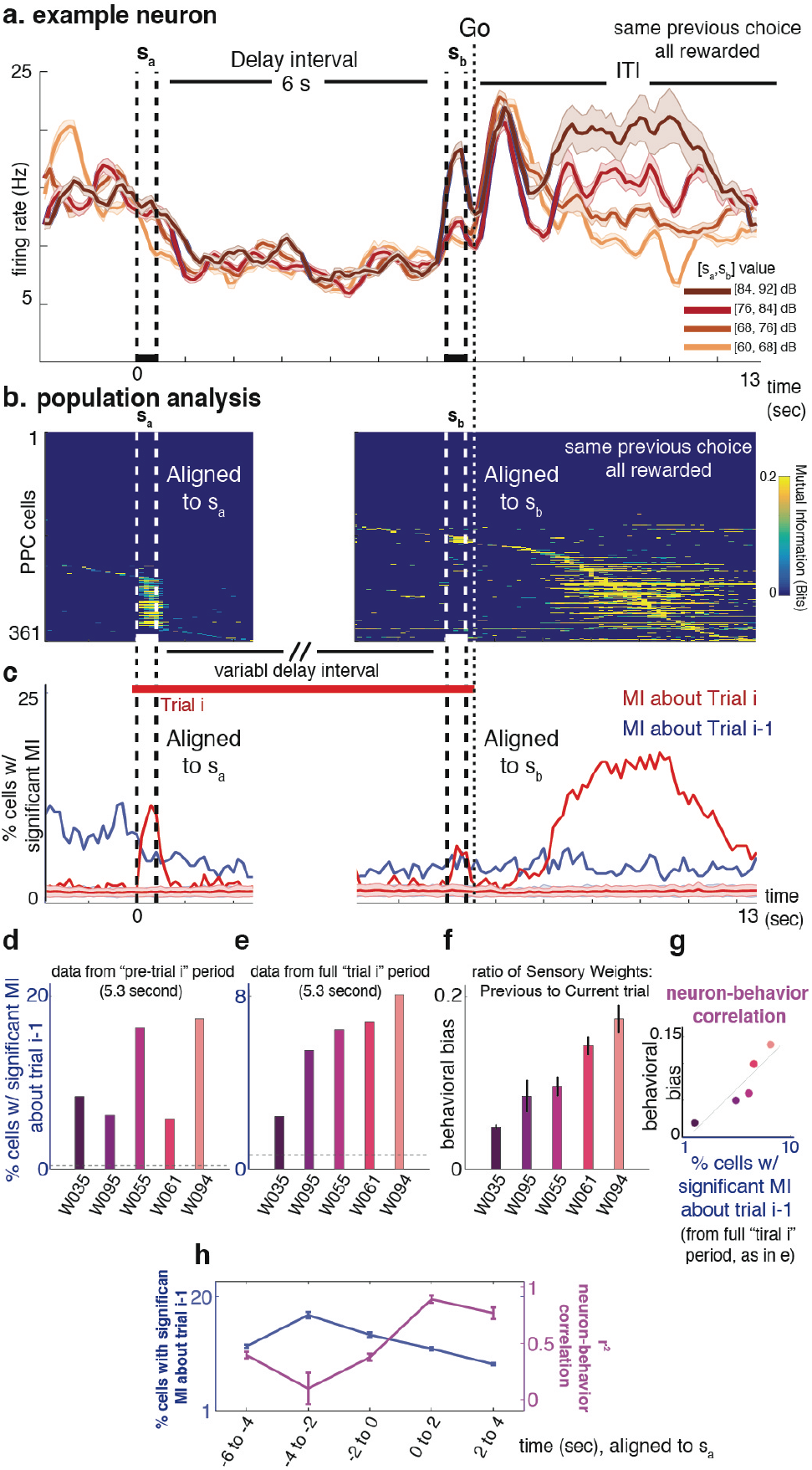
PPC neurons carry more information about previous trial sensory stimuli than about current trial stimuli, which predicts behavioral bias. **a**, Firing rate of example neuron in response to different s_a_ values. For clarity, shows only trials in which animal responded left after Go cue, was rewarded, and delay interval was 6 s. **b**, Mutual information (MI) between each neuron’s firing rate and stimulus pair [s_a_,s_b_], as a function of time. Data is from trials with delay intervals 2-6 s. Left, aligned to start of s_a_; right to start of s_b_. Only MI values significantly larger than shuffled distribution (p < 0.005) are included; non-significant values, dark blue. To control for reward and choice, MI values were calculated using only trials with fixed choice and reward (Methods) **c**, Summary of population analysis, showing % cells with significant coding of stimuli from trial i (red), or trial i-1 (blue). **d**, % cells with significant MI about previous trial, for each of five recorded animals, calculated over 5.3 seconds during ITI before new trial. Horizontal dashed line, % of cells expected by chance from shuffled data (Methods). **e**, As in d, data from 5.3 seconds of new trial. **f**, Behavioral bias for individual animals, calculated as relative sensory history weights (to weights for s_a_ and s_b_ regressors of trial i) from best model fit. **g**, Data from (f) plotted against data from (e), Spearman’s rank correlation r = 1.0, p<0.01, n = 5 rats. **h**. Calculation as in (d-g), using 2 s windows. High neuron-behavior correlation appears concurrently with new trial’s s_a_. Error bars show STD of mean over n = 1000 bootstrap samples with replacement.

Parametric working memory tasks, with their quantifiable behavior, are well-suited to investigating the effect of sensory history on perception and behavior. Rodent versions of these tasks, with semi-automated training, are an efficient platform for causal and cellular-resolution investigation of neural mechanisms. Using this platform, we identified the PPC as an essential node in both the representation and causal effects of sensory stimulus history. This opens a critical window towards a cellular-resolution understanding of long-standing questions about how sensory stimulus history affects working memory and perception. Important issues that can now be addressed include how history representations in PPC interact with current stimulus representations to modulate perception, how history information reaches PPC, and which brain regions connected to PPC are also essential nodes of the circuit.

## Author contributions

AA and CDB conceived the project. AA carried out all experiments and analyzed the data, with the optogenetic inactivations carried out with assistance from CDK. AA gathered Human tactile data fin MED’s laboratory. AA and CDB wrote the manuscript, based on a first draft by AA, with extensive comments by CDK and MED.

## Acknowledgments

We thank C. Duan, R. Low, A. Piet, L. Pinto, and B. Scott, I. Witten for their comments on the manuscript. We thank K. Osorio and J. Teran for animal and laboratory support.

## Methods

### Rat Subjects

A total of 33 male Long-Evans rats (*Rattus norvegicus*) between the ages of 6 and 24 months were used for this study. Of these, 25 were used for behavioral assessments, 6 rats were used for neural recordings, and 7 for optogenetic inactivations. All statistical tests were made between groups with similar sample sizes. Investigators were not blinded to experimental groups during data collection or analysis. Animal use procedures were approved by the Princeton University Institutional Animal Care and Use Committee and carried out in accordance with National Institutes of Health standards.

### Human Subjects (auditory)

11 human subjects (8 males and 3 females, ages 22-40) were tested and all gave their informed consent. Participants were paid to be part of the study and were naive to the main conclusions of the study. The consent procedure and the rest of the protocol were approved by the Princeton University Institutional Review Board.

### Human Subjects (tactile)

14 human subjects (8 males and 6 females, ages 22–35) were tested. Protocols conformed to international norms and were approved by the Ethics Committee of the International School for Advanced Studies (Trieste, Italy). Subjects signed informed consent.

### Rat Behavior

We developed a computerized protocol to train rats, in high-throughput facilities, to perform an auditory delayed comparison task, adapted from a tactile version^6^. All training happens in three-port operant conditioning chambers, where ports are arranged side-by-side along one wall, and with two speakers, placed above the right and left nose ports. **Fig. 1a** shows the task structure. A visible LED in the center port signals the availability of each trial. Rat subjects initiate a trial by inserting their nose into the center port that causes the center LED to turn off. Rats must keep their nose in the center port (“fixation” period) until an auditory “go” cue, a 6kHz pure tone, signals the end of fixation, for 200 ms. Only after the “go” cue, subjects can withdraw and orient to one of the side pokes in order to receive water reward. During the fixation period, two auditory stimuli, ‘s_a_’ and ‘s_b_’, separated by a variable delay, are played for 400 ms. There are short delay periods of 250 ms inserted before ‘s_a_’ and after ‘s_b_’. Stimuli consist of broadband noise (2k-20k Hz), generated as a series of Sound Pressure Level (SPL) values sampled from a zero-mean normal distribution. Overall mean intensity of sounds vary from 60-92 dB. Rats should judge which of s_a_ and s_b_ had the greater SPL standard deviation. If s_a_ > s_b_ then the correct action is to poke in the right side poke in order to collect the reward, and if s_a_ < s_b_ rats have to orient to the left side poke. Trial durations are independently varied on a trial-by-trial basis, by varying the delay interval between the two stimuli, that can be as short as 2 s or as long as 12 s. Rats progressed through a series of shaping stages, before the final version of the delayed comparison task, in which they learned to 1) associate light in the center poke with availability of trials, 2) associate special sounds from the side pokes with reward, 3) maintain their nose in the center poke until they hear an auditory “go” signal, and 4) compare the two s_a_ and s_b_ stimuli.

Although substantial amount of data has been collected on all delay intervals from 2 to 12 seconds, in this manuscript we focus on delay durations of 2, 4 and 6 seconds, since most of the rats were consistently trained on these values. Training began when rats were 2 months old and typically required #3# to #4# months for rats to display stable performance on the complete version of the task.

### Human Auditory Behavior

Similar auditory stimuli to those used for rats were used in the human version of the task. In this experiment, subjects received, on each trial, a pair of sounds played from ear surrounding noise canceling headphones (brand 233621-H501). The subject self initiated each trial by pressing the space bar on the keyboard. The first sound was then presented together with a green square on the left side of a computer monitor in front of the subject. This was followed by a delay period, indicated by “WAIT!” on the screen, then the second sound was presented together with a red square on the right side of the screen. At the end of the second stimulus and after the go cue, subjects were required to compare the two sounds and decide which one was louder, then indicate their choice by pressing the “k” key with their right hand (second was louder) or the “s” key with their left hand (first was louder). Written feedback about the correctness of their response was provided on the screen, for individual trials, as well as the average performance updated every 10 trials.

### Human Tactile Behavior

In a separate set of experiments, run at International School For Advanced Studies (SISSA), human subjects performed the tactile version of the task. The details of this task have been previously described and the behavior has been characterized ^6^. Briefly, at each trial two noisy vibration stimuli, interleaved with a variable delay interval, were delivered to the subject’s fingertip on their ## (left?) hand. Subjects viewed a computer monitor and wore headphones that presented acoustic noise and eliminated ambient sounds. To start a trial, the subject pressed the keyboard up arrow with their right hand. This triggered presentation of the two stimuli. After a post-stimulus delay, a blue panel was illuminated on the monitor, and the subject pressed the left or right arrow on the keyboard, signifying selection of the first or the second stimulus as greater, respectively. They received feedback (correct/incorrect) on each trial via the monitor. Human experiments were controlled using LabVIEW software (National Instruments, Austin, Texas).

### Stimulus Set

If the first stimulus, s_a_ was fixed across all trials and only the second stimulus s_b_ changed, subjects might solve the task by ignoring the first stimulus and applying a constant threshold to the second stimulus. Likewise, if the second stimulus was fixed, subjects might apply a constant threshold on the first stimulus. To prevent such alternative strategies, it is necessary to vary both sa and sb, and use a set of stimuli composed of pairs of s_a_ and s_b_ which guarantees that across trials the same value of SPL standard deviation is randomly presented for the first stimulus or the second stimulus. Stimuli along the diagonal in Fig. 1b represent such a stimulus set. A minimum of 8 pairs of stimuli span a wide range of SPL standard deviation values (**Fig. 1b**). Using this stimulus set, if the subject were to ignore either of “s_a_” or “s_b_”, then the maximum performance would be 63%. The mean amplitudes of stimuli were evenly distributed in a logarithmic scale (linear in dB). The diagonal line represents s_a_ = s_b_; all of the stimulus pairs on one side of the diagonal are associated with the same action, and all have the same ratio of s_a_ to s_b_. For each trial, one of these 8 pairs of stimuli is randomly selected to determine s_a_ and s_b_.

### Psychometric curves

In some sessions, we used stimulus pairs that were closely spaced along **s_a_** (**Fig. 1b**) to measure the psychometric discrimination threshold. Psychometric plots (as shown in **Fig. 1e** and **Fig. 3c**) show the probability of the subject responding leftward as a function of the difference between **s_a_ and s_b_** when **s_b_** is fixed. The fits were to a 4-parameter logistic function of the form

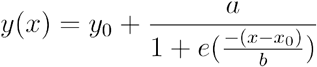

Where *y*_0_ is the left endpoint, *y*_0_ + *a* is the right endpoint, *x_0_* is the bias, and *a*/4*b* is the slope. Fits were done using the non-linear least square regression method (nlinfit.m function) in Matlab2013.

### Regression Model of Behavior

Our semi-automated training protocol facilitated the generation of a behavioral data set comprising 468,165 trials from 25 trained animals, which in turn enabled statistical characterization of the decision-making process. In order to quantify the rats’ behavior we carried out an analysis to “weigh” the contributions of s_a_ and s_b_ on the current trial and several trials in the past, as well as the history of choice and reward on the animal’s choice in the current trial. Using the data generated by concatinating multiple training sessions, we fit the animal’s choice with a logistic regression model that allows the linear combinations of s_a_ and s_b_ and other desired factors. The linear combination is then mapped nonlinearly into the animal’s choice, i.e. the probability of trials in which the subject judged s_a_ > s_b_, through a logistic function as:

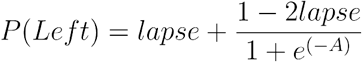

Where

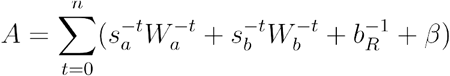

Where 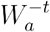 and 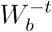 are coefficients of the s_a_ and s_b_ regressors, respectively, from *t* trials back. 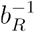 is the “correct side” on the previous trial: left = +1, right = −1. This regressor captures the win-stay/lose-switch strategy. *β* is the baseline regressor that captures the overall (stimulus-independent) bias of the subject in calling s_a_ > s_b_ (for instance, a bias against turning right, the side associated with the judgment of s_a_ > s_b_). The absolute values of all of the regressors are normalized between 0 and 1. We used the log-likelihood as the cost function C:

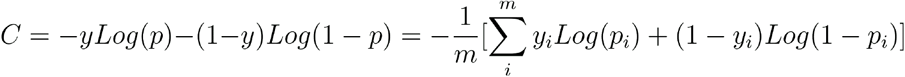

The model was fit using a gradient descent algorithm to minimize the negative log-likelihood cost function. We used the “sqp” algorithm in the fmincon function from Matlab 2013. Weights were calculated using L2- regularization to prevent overfitting. The hyperparameter value (lambda) was selected independently for each rat using evidence optimization, based on 5-fold cross-validation. Different variants of the model, that systematically study the relevance of various sensory and reward history factors^18,19,32^, capturing not only win-stay/lose-switch, but also perseveration, are discussed in the **Extended Data Fig. 6**.

### Model comparison and cross-validation

All models were fit separately for each individual rat (n = 25), using 200 runs of 5-fold cross-validation. For each run we calculated the log-likelihood of the test dataset given the best-fit parameters on the training set (Logl). We also calculated the log-likelihood of the test dataset for the mean value of %Left (the experimentally-measured fraction of trials in which the animal went left). This gives us a null log-likelihood reference value (Logl_0_). In order to quantify the efficiency of each model we defined the “cross-validated bit/trial” (CV-bit/trial) as the trial-averaged excess likelihood of the model compared to the null model^33^:

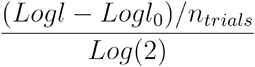

For each model, we first chose the optimal regularization value (lambda) that would maximize the CV-bit/trial. To compare different models we calculated the median value of CV-bit/trial across 10000 fits for each subject. Since in this method we measure the log-likelihood using the cross-validated data, it automatically takes care of the overfitting problem, such that if additional parameters of one model result in overfitting in the training set, it would penalize it in the cross-validated test set.

### Optogenetic virus injection and fibre implantation

For optogenetic perturbation experiments, the general surgery techniques and fiber etching follow previous reports ^34^, except that we began the construction with a standard off the shelf 50/125 μm LC-LC duplex fibre cable (http://www.fibercables.com), instead of the usual FC-FC duplex fiber cables. The cable jacket, strengthening fibres, and outer plastic coating (typically white or orange) were fully removed leaving 1 cm of the fibre optic cable and the inner plastic coating (typically clear) intact. Then 2 mm of the fibre tip (with final layer of plastic coating still attached) was submerged in 48% hydrofluoric acid topped with mineral oil for 85 min, followed by water for 5 min (submerging 5 mm), and acetone for 2 min (to soften the plastic). The plastic coating was then gently cut with a razor and pulled off with tweezers to reveal a 1 mm sharp-etched fibre tip. Enough plastic was removed, depending on the depth of the targeted site, to ensure that only the glass fibre optic would be inserted into the brain.

For viral injections, 2 μl of adeno-associated virus (AAV) (AAV5-CaMKIIα-eNpHR3.0-eYFP), that drives expression of the light-activated inhibitory opsin halorhodopsin eNpHR3.0, under the CaMKIIα promoter, coupled to eYFP, was lightly dyed with fast green powder and front loaded into a glass pipette mounted to a Nanoject (Drummond Scientific) prefilled with mineral oil. The pipette tip was manually cut to ~30 μm diameter. Five closely spaced injection tracts were used for each animal. For the central injection tract (AP -3.8 mm and ML 2.5 mm from Bregma, Fig. 3a, brain image from Paxinos and Watson^31^), one injection of 23 nl were made every 100 μm in depth starting 100 μm below the brain surface of the PPC for 1.5 mm. Four additional injection tracts were completed, using procedures identical to the central tract, one 500 μm anterior, posterior, medial, and lateral from the central tract. Each injection was followed by a 10 s pause, with 1 min following the final injection in a tract before the pipette was removed. A total of 1.5 μl of virus was injected over a 30-min period consisting of ~160 separate injections. A chemically sharpened fibre optic (50 μm core, 125 μm cladding) was then lowered down the central injection tract to a depth of 1 mm. The craniotomy was filled with kwik-sil (World Precision Instruments), allowed to set for 10 min, and the fibre optic was secured to the skull with C&B Metabond and dental acrylic. Dental acrylic covered all of the incision site and allowed only the LC connector to protrude. Halorhodopsin expression was allowed to develop for 6 weeks before the behavioural testing began.

### Optogenetic perturbation

The animal’s implant was connected to a 1-m patch cable attached to a single fibre rotary joint (Princetel) mounted on the ceiling of the behavioural chamber. This was connected to a 200 mW, 532 nm laser (OEM Laser Systems) operating at 25 mW, which was triggered with a 5 V transistor-transistor logic (TTL) pulse. Laser illumination occurred on 20% of randomly selected trials. See **Extended Data Fig. 7** for physiological confirmation of optogenetic inactivation effects in an anesthetized animal. Based on our previous quantifications of optogenetic effects ^34^, we estimate that using eNpHR3.0 we can inhibit, almost entirely, neurons in a radius ~750μm from the tip of the optic fibre, amounting to an sphere of ~1500μm in diameter.

### Recordings

6 animals were implanted with microwire arrays in their left or right PPC (n=3 in rPPC, n=3 in lPPC, see **Extended Data Fig. 7** for histological localization of electrodes). The target region was accessed by craniotomy, using standard stereotaxic technique (centered 3.8 mm posterior to the bregma and 2.5 mm lateral to the midline). Dura mater was removed over the entire craniotomy with a small syringe needle. The remaining pia mater, even if usually not considered to be resistant to penetration, nevertheless presents a barrier to the entry of the microelectrode arrays, due to the high-density arrangement of electrodes in the multi-channel electrode arrays. This dimpling phenomenon, when the electrodes are pushing the brain cortex down without penetrating, is more pronounced for arrays with larger number of electrodes. In addition to potentially injuring the brain tissue, dimpling is a source of error in the determination of depth measurements. Ideally, if dimpling could be eliminated, the electrodes would move in relation to the pial surface, allowing effective and accurate electrode placement. To overcome the dimpling problem, we implemented the following procedure. After the craniotomy was made, and the dura was carefully removed over the entire craniotomy, a petroleum-based ointment such as bacitracin ointment or sterile petroleum jelly (Puralube Vet Ointment) was applied to the exact site of electrode implantation. The cyanoacrylate adhesive (Vetbond Tissue Adhesive) was then applied to the zone of the pia surrounding the penetration area. This procedure fastens the pia mater to the overlying bone and the resulting surface tension prevents the brain from compressing under the advancing electrodes. Once the polymerization of cyanoacrylate adhesive happened, over a period of few minutes, the petroleum ointment at the target site was removed, and the 32 electrode microwire array (Tucker-Davis Technologies (TDT), Alachua FL) was inserted by slowly advancing a Narishige hydraulic micromanipulator. After inserting the array(s), the remaining exposed cortex was covered with biocompatible silicone (kwik-sil, World Precision Instruments), and the microwire array was secured to the skull with C&B Metabond and dental acrylic.

During the 10 days of recovery time, rats had unlimited access to water and food. Recording sessions in the apparatus began thereafter, using Neuralynx acquisition systems. Extracellular activity of the PPC neurons was manually sorted into single units and multi-units, based on the spike waveform and the refractory period observed in the interspike interval histogram, using SpikeSort3D software. In total 1081 single or multi-units were recorded in the PPC of 6 rats. Only neurons whose overall mean firing rate within the session was at least 2 Hz were included in the analysis, for a total of 456.

## Neural Analysis

### Mutual Information

In order to quantify the type and amount of information that the PPC neurons carry about various task parameters we computed Shannon’s Mutual Information ^35^. In this formulation, the amount of information which can be extracted from the firing rate of a neuron R, about the task-related parameter X can be computed as:

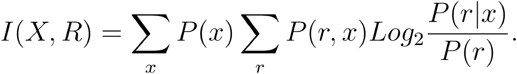

Where *P(r|x)* is the conditional probability of observing a neuronal response r given the presentation of the task parameter x, *P(r)* is the marginal probability of occurrence of neuronal response r among all possible responses, and *P(x)* is the probability of task parameter x. Information measured in this way quantifies how well an ideal observer can discriminate between members of a stimulus set based on the neuronal responses of a single trial 36. For each trial, neuronal response was defined as the rate of spiking during a time windows of 100 ms. The conditional probability in the above formula is not known *a priori* and must be estimated empirically from a limited number, N, of experimental trials for each stimulus. Limited sampling of response probabilities can lead to an upward bias in the estimate of mutual information ^37^. In order to correct for this bias, we used a combination of two techniques. First we estimated and corrected the bias based on the Quadratic Extrapolation (QE) method ^38^, that assumes the bias can be accurately approximated as second order expansions in 1/N. Then we used bootstrap procedure (shuffling) that consists of many rounds of pairing stimuli and responses at random in order to destroy all the information that the responses carry about the stimulus. Due to limited data sampling, the information computed using the bootstrapped responses may still be positive. The average value of the bootstrapped information was then used to estimate the residual bias of the information calculation, and was subtracted out. Moreover, the distribution of bootstrapped information values were used to build a non-parametric test of whether the corrected information computed using QE method is significantly different from zero ^39^.

Using the MI distribution from a shuffled dataset, at each time bin, where trials are randomly labeled, we first calculated the bin-by-bin estimate of the percent of cells with significant value of MI expected by chance (Fig. 4c, shuffled data). We then computed the average over ITI or “trial i” duration to find the mean values depicted by dashed lines in Fig. 4d-e. To control for reward and choice, MI values were calculated using only trials with fixed choice and reward, and only then averaged across different, separately calculated reward and choice groups.

### Code and data availability

All software used for behavioral training is available on the Brody lab website at http://brodylab.org/auditory-pwm-task-code. Software used for data analysis, as well as raw and processed data, are available from the authors upon reasonable request.

---

**Extended Data Fig. 1.**
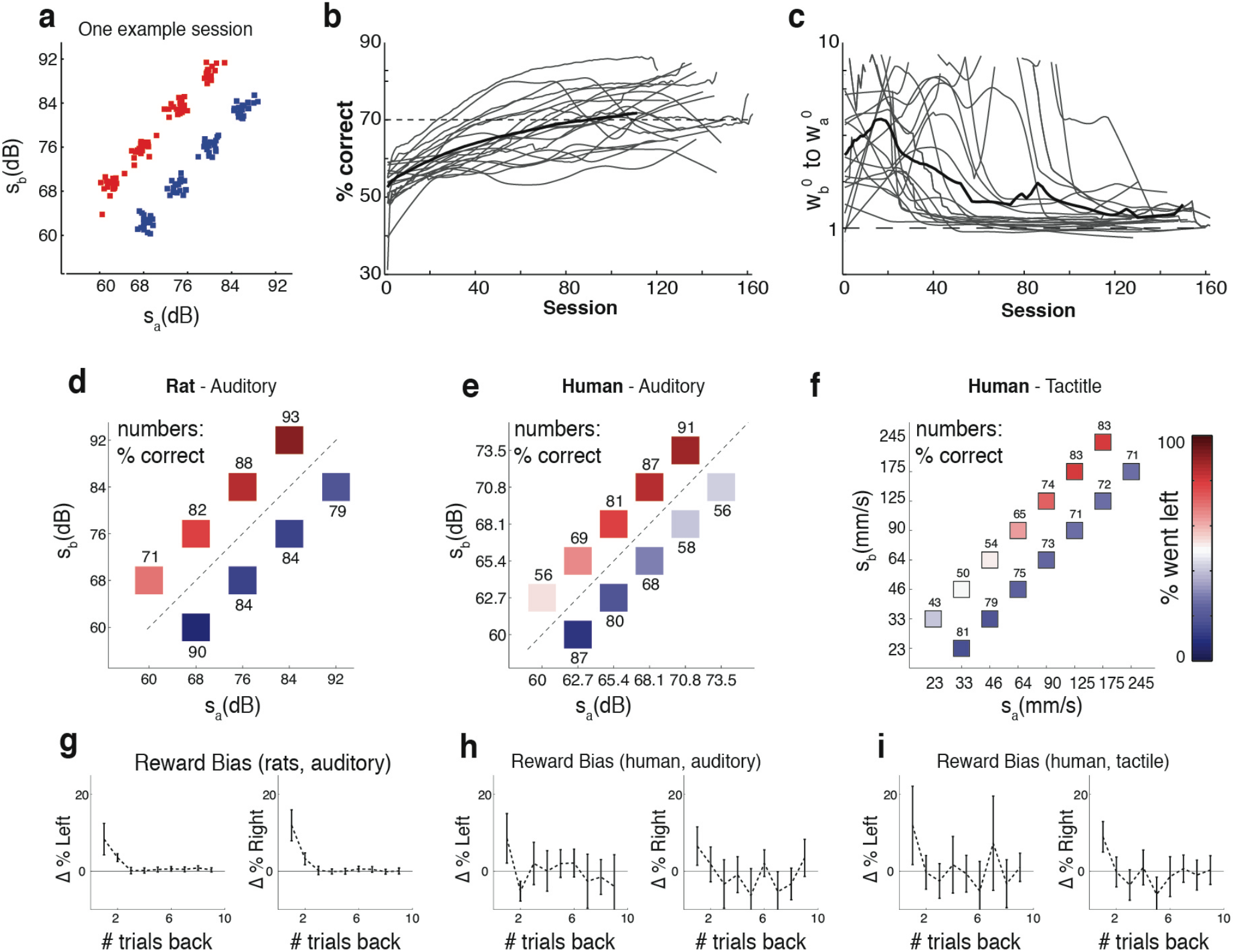
Full stimulus set - Learning curve - Mean performance - Reward bias. **a**, Each stimulus is made of a series of Sound Pressure Level (SPL) values sampled from a zero-mean normal distribution, and standard-deviation value of s. For each trial, SPL values are randomly drawn and therefore due to sampling statistics, the actual standard deviation value of the stimulus always differed slightly from its designated value. The coordinates of each small box represent the actual joint values of [s_a_, s_b_] for one sample training session. **b**, Individual gray lines shows learning curves presented as the change in % correct over months of training, for n = 25 rats. Average rat (black line) reaches 70% of performance after 90 sessions. **c**, Learning curve presented as the ratio of the best fit weights for the second stimulus, sb, to the first stimulus, sa, using the model described in Figure 2e (3-parameter, “No history” version). **d**, Rat auditory working memory performance, data from 21 rat subjects (total of 468,165 trials) are separated by [sa,sb] pair but averaged across subjects and over different delay durations (2–8 sec). **e**, Human auditory working memory performance. For human, interstimulus delay varied randomly from 2 s to 6 s. (11 subjects, 12623 trials). **f**, Human tactile working memory performance. Similar to panel “e” but for humans engaged in the tactile version of the task. In this task, interstimulus delay varied randomly from 2 s to 8 s. Data from 14 human subjects (total of 4694 trials) are pooled together. **g**, Reward history bias. Left panel: y-axis shows, for “turn left” trials and as a function of k, % went left when the k^th^ trial back was rewarded on the left, minus % went left when the k^th^ trial back was rewarded on the right. Right panel, complementary plot for “turn right” trials: % went right when the k^th^ trial back was rewarded on the right, minus % went right when the k^th^ trial back was rewarded on the left. Data from n = 21 rats. Each points show the mean value of the bias over subjects. Errorbars show 95% confidence interval. **h-i**, similar to (g) for human auditory (h, n = 11 subjects) and tactile (i, n = 14 subjects) PWM task.

**Extended Data Fig. 2.**
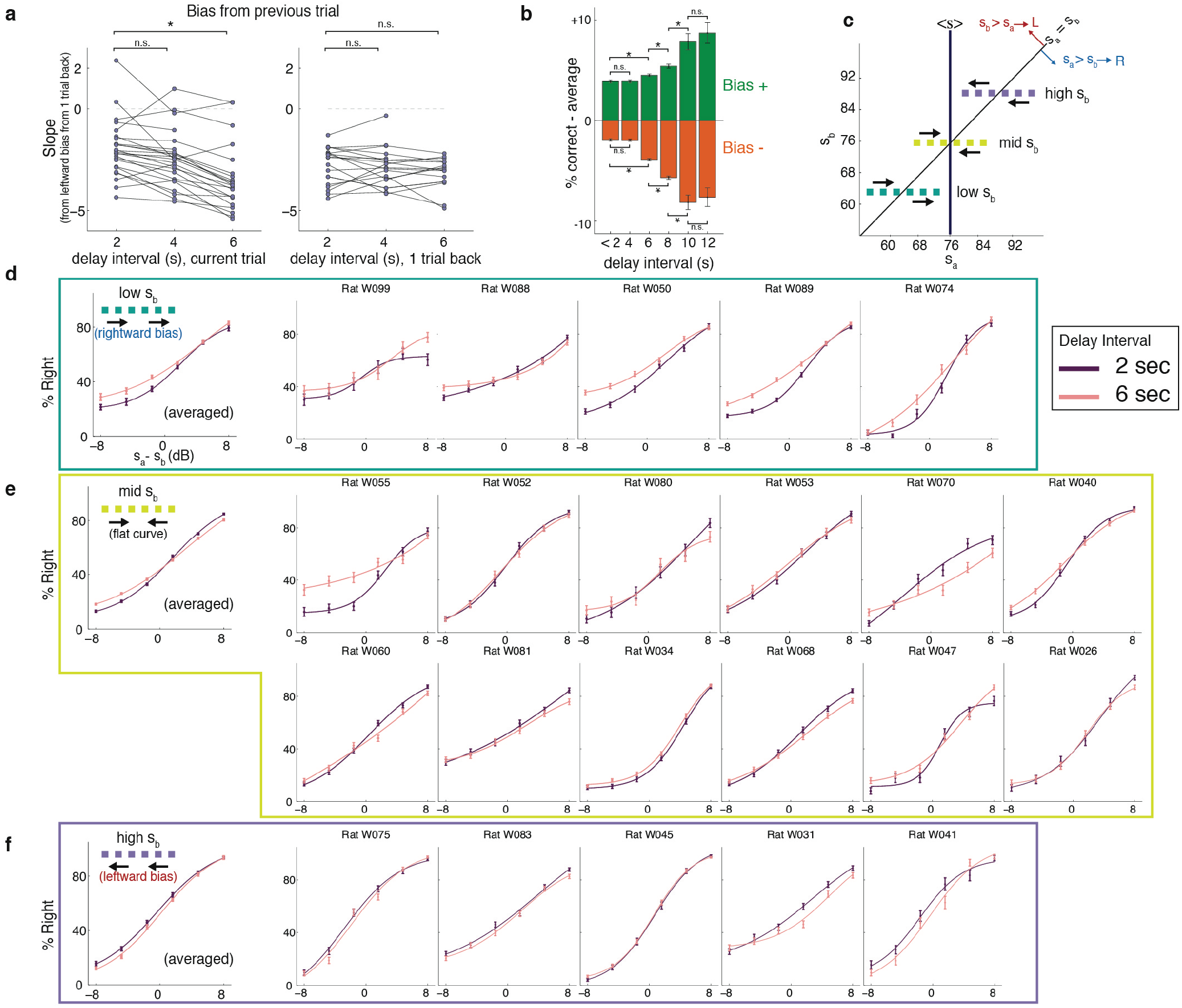
Contraction bias grows as a function of the current trial’s WM delay interval. **a**, Slopes from linear fits to %leftward bias (as in **Fig. 2a**), for rats that were each trained on delay intervals of 2, 4, and 6 sec (n=21). Left shows behavioral bias (“% went left - average”) as a function of WM delay interval of the current trial. Right shows behavioral bias as a function of WM delay interval from 1 trial back. Each dot represents a rat; lines connect the different delay intervals for each rat. Left: from a one-sided paired t-test, 2 vs. 4 second: p = 0.012, 2 vs. 6 second: p < 0.001; 4 vs 6 second: p = < 0.001. ‘*’ indicates significant difference (p < 0.001, one-sided paired t-test). Right: 2 vs. 4 second: p=0.76, 2 vs. 6 second: p = 0.37; 4 vs 6 second: p = 0.65. The behavioral bias grows with for greater current WM delay period, but no significant dependence on the previous trial’s WM delay period is found^26^. **b**, Percent correct averaged across all Bias+ trials or all Bias− trials, relative to overall average performance, as a function of WM delay interval on the current trial. Data is pooled from a dataset in which different rats were trained on different sets of delay intervals; data for each delay interval may thus contain different rats than data for other delay intervals (n = 25 rats total). Errorbars show standard deviation of mean. As in panel (a), behavioral effect grows as a function of the current WM delay period. ‘*’ indicates significant difference (p < 0.001, one-sided t-test). **c**, Schematics of stimuli used for three different psychometric curves: “high s_b_”, in which contraction bias would lead all the s_a_ stimuli to be treated as lower than they actually were (indicated by the leftward arrows), producing a rightward shift of the psychometric curve; “mid s_b_”, in which contraction bias would lead all the s_a_ stimuli to be treated as closer to s_b_ than they actually were, producing a flattening of the psychometric curve; and “low s_b_”, in which contraction bias would lead all the sa stimuli to be treated as higher than they actually were, producing a leftward shift of the psychometric curve. **d**, Psychometric curves for “high s_b_” trials, averaged across animals and separately for each individal animal, for trials with a 2 sec WM delay interval, and for trials with a 6 sec WM delay interval. Curves are fits to a 4-parameter logistic function (see Methods). As the WM delay interval grows, the leftward shift predicted by contraction bias shift is more pronounced. For each individual animal n = 120 sessions of data were used. Errorbars show standard error of mean over sessions. **e**, as in panel d, but for the “mid s_b_” trials. As the WM delay interval grows, the flattening predicted by contraction bias is more pronounced. **f**, as in panel d, but for the “low s_b_” trials. As the WM delay interval grows, the rightward shift predicted by contraction bias is more pronounced.

**Extended Data Fig. 3.**
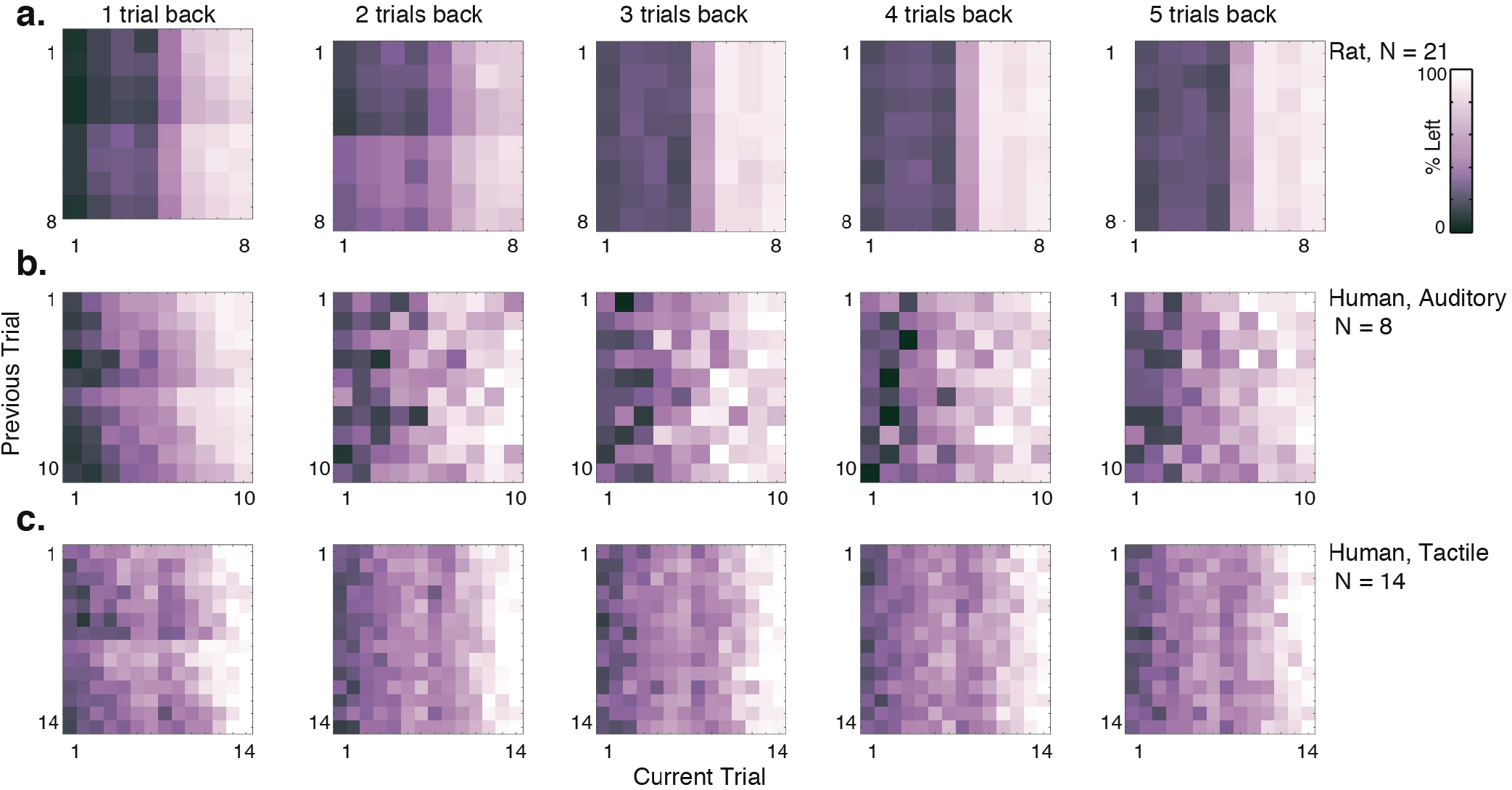
Sensory History matrix, from 1 to 5 trials back. **a**, Stimulus history matrix, as described in Figure 2b, when %left is shown given any combination of the stimuli in current trial (x-axis) and n-trials back (y-axis), n = 1,2,3,4,5. Trial numbers indicate pairs of [s_a_ s_b_]: 1=[68(dB) 60(dB)], 2=[76 68], 3=[84 76], 4=[92 84], 5=[60 68], 6=[68 76], 7=[76 84], 8=[92 84]. Data from N = 21 rats, composing total of 381,612 trials is used in this analysis. Trial numbers are: 1=[62.7(dB) 60(dB)], 2=[65.4 62.7], 3=[68.1 65.4], 4=[70.8 68.1], 5=[73.5 70.8], 6=[60 62.7], 7=[62.7 65.4], 8=[65.4 68.1], 9=[68.1 70.8], 10=[70.8 73.5]. **b**, Similar to panel “a”, for human auditory task. **c**, Similar to panel “a”, for human tactile task. Trial numbers are: 1=[33(mm/s) 23(mm/s)], 2=[46 33], 3=[64 46], 4=[90 64], 5=[125 90], 6=[175 125], 7=[245 175], 8=[23 33], 9=[33 46], 10=[46 64], 11=[64 90], 12=[90 125], 13=[125 175], 14=[245 175].

**Extended Data Fig. 4.**
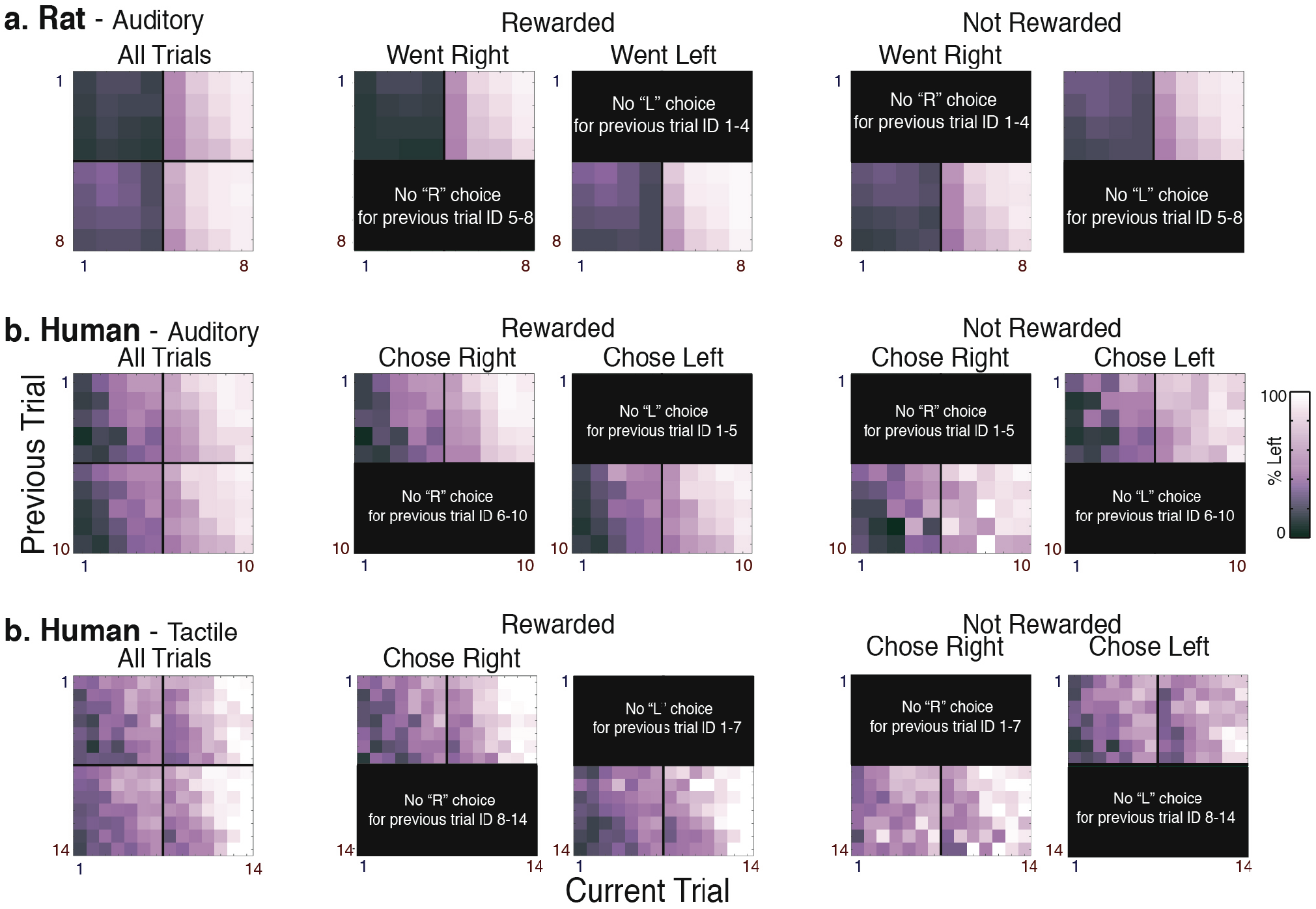
Sensory History matrix, controlled for Reward and Choice. Similar to Extended Figure 2, except that in this plot only trials for which the previous trial was a “turn right” trial and the animal was rewarded are included. Therefore, modulation by previous trial cannot be due to “action” or “reward” history. Trial numbers are similar to **Extended Data Fig. 3**.

**Extended Data Fig. 5.**
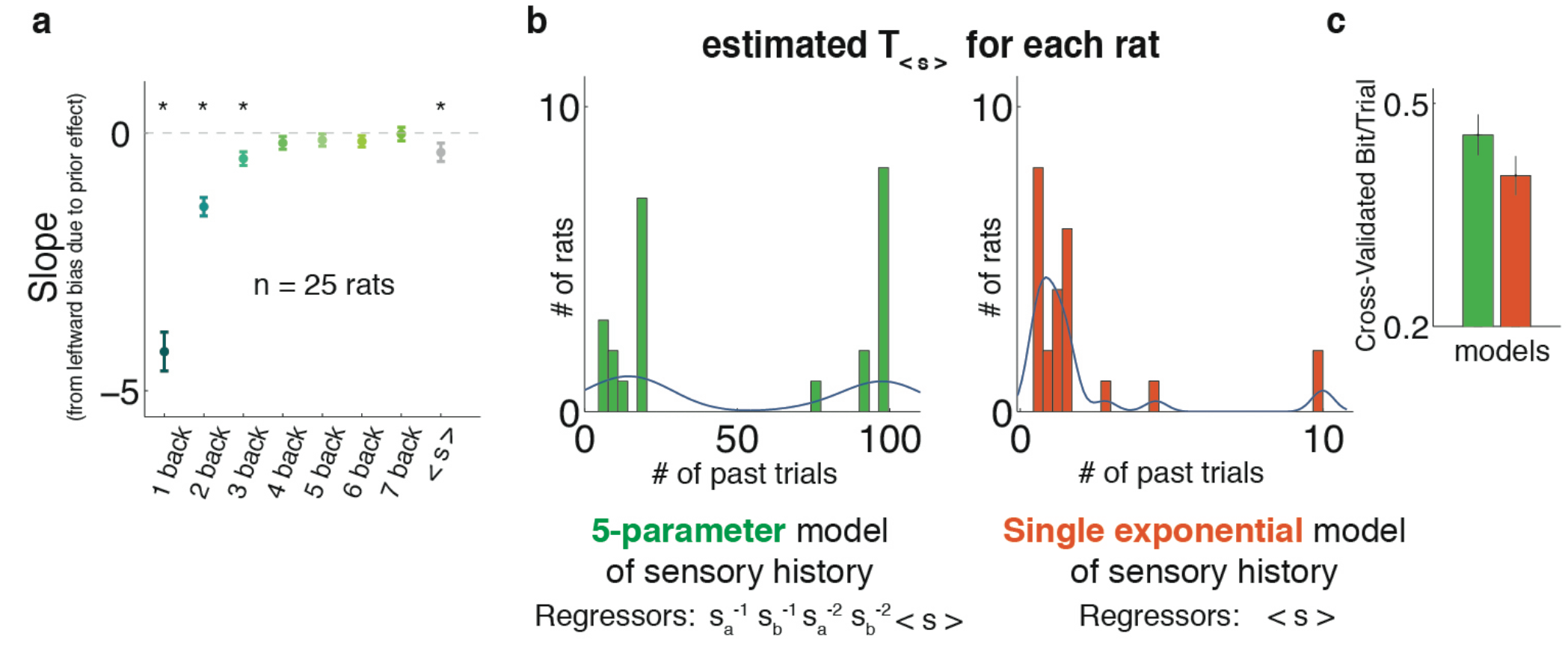
Estimating optimal window of < s >. **a**, Slopes from linear fit to the %leftward bias from n-back trials (n=1:7, as in **Fig. 2a** where n=1 was used), and also < s > which is a window of 17 trials, from n=4 to n=20, in gray. Each point shows mean of the slope values over n = 25 rats. Errorbars show 95% confidence intervals. **b**, For each rat the optimal exponential window over the past trials was estimated such that it would maximize the cross-validation bit/trial measurement. Two models are compared here: green shows the distribution of taus from a model that has 5 regressors to account for the sensory history - first and second stimulus from the 2 trials back and a separate exponential window over the remaining past trials (Fig 2d). In orange, instead, only one regressor which is a single exponential window over all the past trials accounts for the sensory history. In the single-exponential model, the best fit value of tau comes out very small, practically as if only past 1 or 2 trials back are inducing most of the effect. **c**, The 5-parameter model of sensory history outperforms the single-exponential model. 200 iterations of 5-fold cross validatation has been used to calculate the “Cross-Validated bit/trial” (Methods). Accordingly each bar shows the mean of n = 1000 datapoints. Errorbars show

**Extended Fig. 6.**
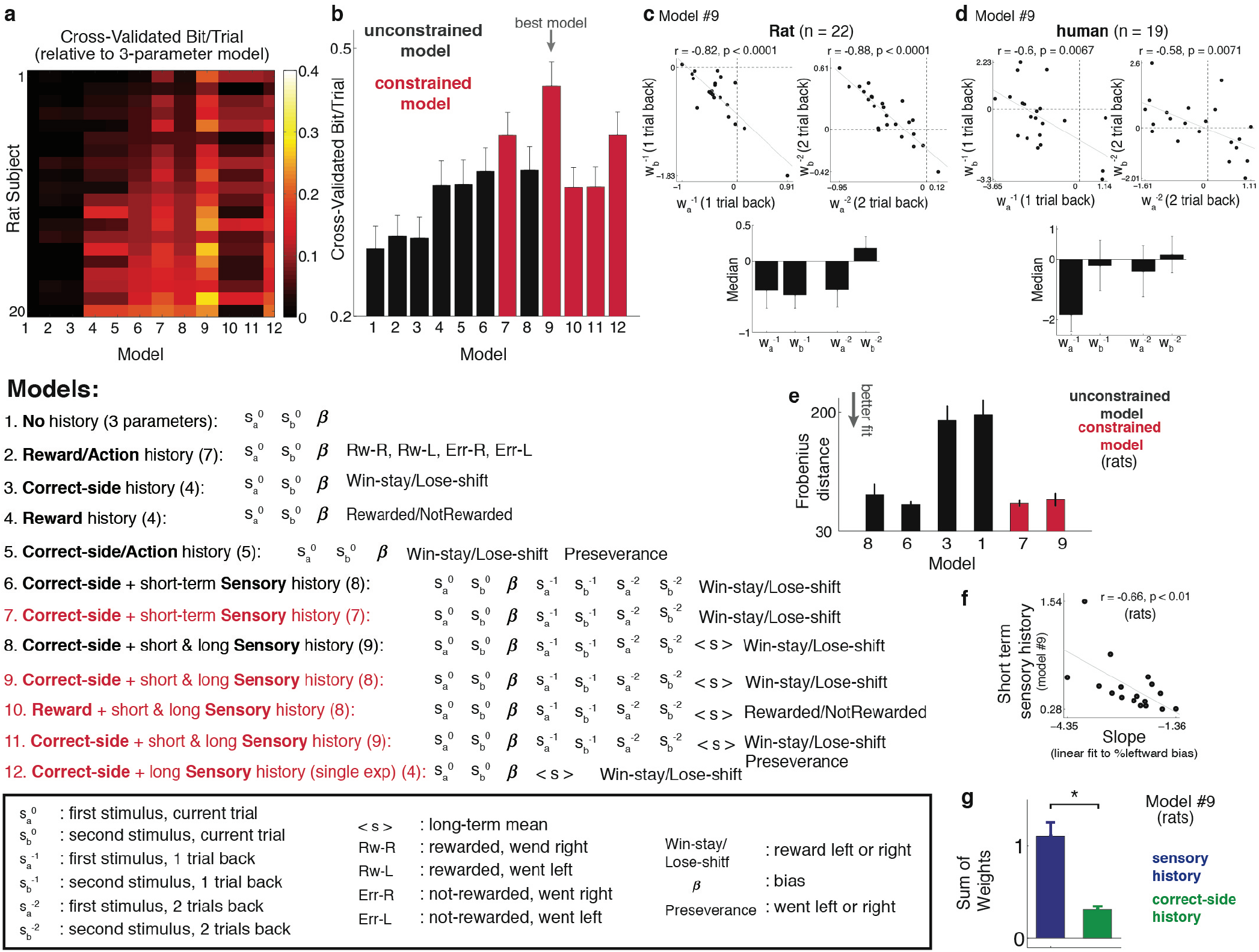
Model comparison. **a**, Model comparisons, 200 runs of 5-fold cross validation were done, on data from each rat, in order to find the best fit parameters and to compare different model fits using the “Cross-Validated Bit/Trial” quantity defined as the relative value of the log likelihood of each model, to the null log likelihood, normalized in log2. Removing one parameter by constraining the regression weights on the current trial’s s_a_ stimulus plus the weights on previous sensory stimuli to add to 1 (constrained model, in red) improved performance on cross-validated data compared to the unconstrained model (in black). **b**, Mean value of “CV Bit/Trial” for different variants of the model as in a, over n = 20 rats. Error bars show SEM. Unconstrained models are in black, whereas red shows constrained models. **c**, Top, raster plots of W^−t^_a_ versus W^−t^_b_ (t = 1,2, from the model #9). Each dot represent a subject. Pearson correlation values (r), and corresponding two-sided p-values are shown for each plot. Bottom, median value of W^−t^_a_ and W^−t^_b_(t = 1,2), across animals. Errorbars show median absolute deviation. **d**, similar to c, for human subjects (auditory and tactile tasks are pooled together). Similar to rat subjects, model #9 gains the best performance for human subjects, as well (data not shown). **e**, to compare the Sensory History Matrix from the real data to the ones predicted from the best model fits. Frobenious distance norm was used, defined as the square root of the sum of the absolute squares of the difference between elements of two matrices. Frobenious distance is a measure of similarity and the smaller the value, the more similar the two matrices. Frobenious distance is calculated separetely for individual rats and each bar shows its mean value over n = 20 rats. Errorbars show SEM. **f**, Scatter plot of “slopes from linear fits to %leftward bias (**Fig. 2a**)” versus “short-term sensory history” i.e. sum of weights for s_a_^-1^, s_b_^-1^, s_a_^-2^ and s_b_^-2^, from model #9. This plot shows significant correlation between the two measurements (Pearson correlation, r = −0.66, two-sided p = 0.0084, n = 17 rats), suggesting that when our logistic fit coefficients are particularly large, the subjects also have a particularly large contraction bias. **g**, examining the weights in the regression model #9, which is determined to be the best model, shows that the weights for sensory history terms are significantly larger than those for the correct-side history term (paired-sample ttest, p < 0.0001, n = 22 rats). Data from individual animals are used to fit the model and bars show the mean value of sensory history weights (in blue), and correct-side history weight (in green), over fit values from n = 22 animals. Errorbars show SEM. Moreover, the sensory history regressor term, i.e. sum of “sensory history weights x regressors” produces larger variance over trials (0.38) compared to the correct-side regressor (0.11), indicating a bigger impact on trial-by-trial behavior.

**Extended Data Fig. 7.**
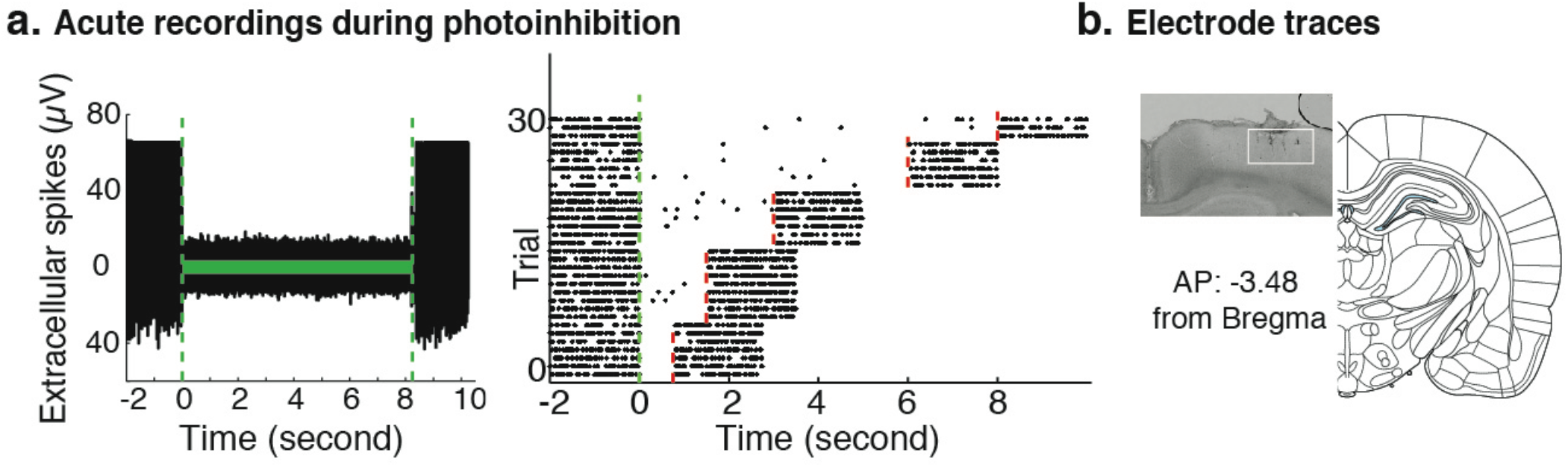
Physiological and histological confirmations. **a**, Physiological confirmation of optogenetic inactivation effect in an anesthetized animal. Left, single trace of acute extracellular activity of an example cell in the PPC, expressing eNpHR3.0, is shown in response to light stimulation. Laser illumination period (8 s) is marked by the light green bar. Right, raster-plot for 32 trials, for variable durations of light stimulation. The green vertical dashed line indicates start of the laser illumination. The laser was on for variable durations of 750ms, 1500ms, 3000ms, 6000ms, or 8000ms. Laser turning off is indicated by the red vertical dashed line. Recordings continued for 2 seconds after the laser was turned off. **b**, histological localization of electrodes targeting PPC. The inset shows example of electrode locations in a coronal slice at AP=3.48 from Bregma. In all cases, the electrode and fiber placements in PPC were within between 2.8 and 4 mm anterior to Bregma and between 2 and 3.5 mm lateral to the midline. Brain image is taken from Paxinos and Watson 2004^31^

**Extended Data Fig. 8.**
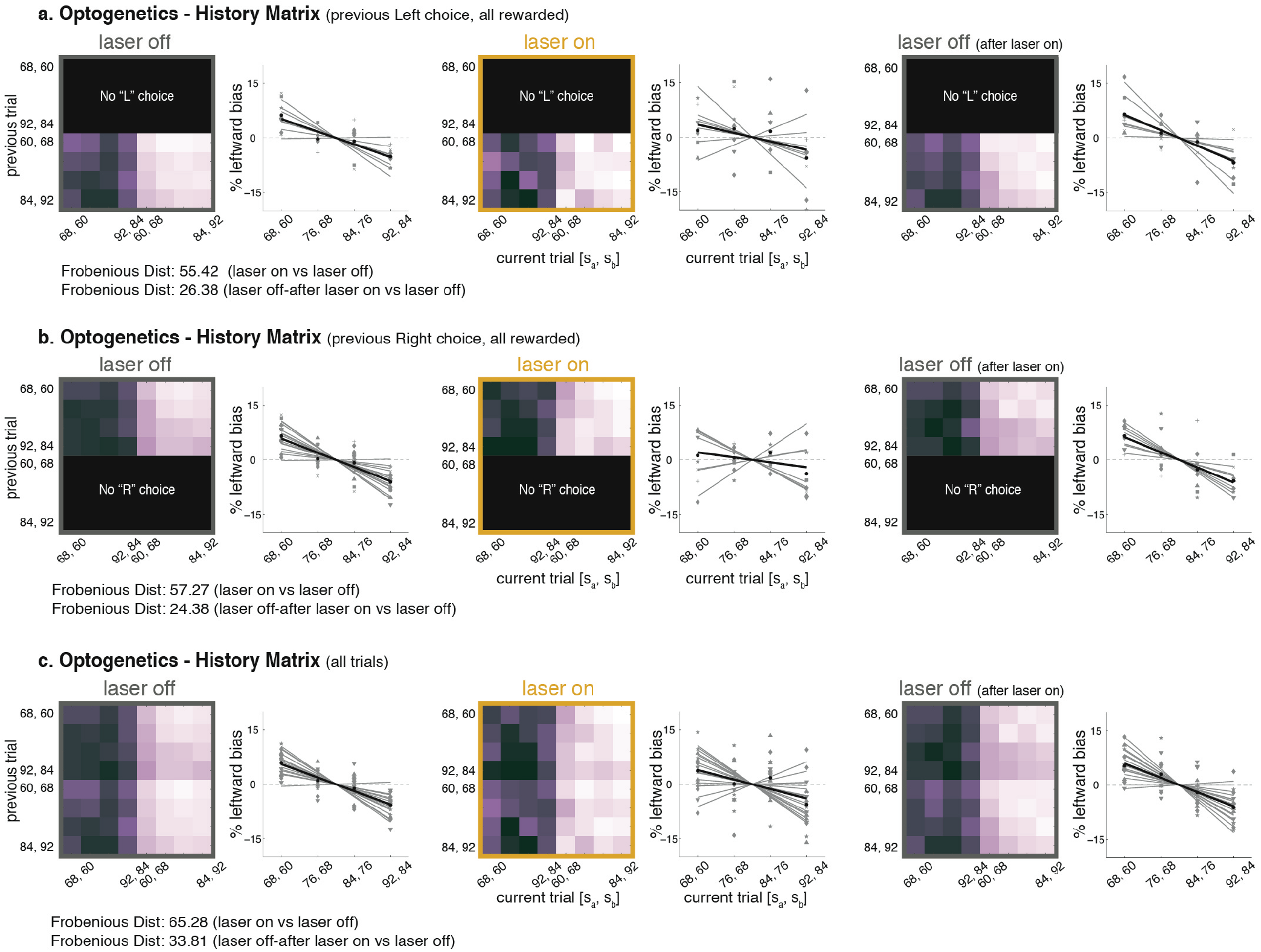
Optogenetics - PPC inhibition reduces leftward bias due to past sensory stimuli. **a**, Sensory History Matrix, and leftward biases due to past sensory stimuli, similar to **Figure 2a-c**, but now for three types of trials: “laser off” trials (two leftmost panels) which consist of trials with no PPC inactivation on either “current” or “previous” trial; “laser on” trials (two middle panels) which consist of trials with PPC inactivation on “current” trial; “laser off after laser on” trials (two rightmost panels) which consist of trials immediately after the “laser on” trials. This last set controls for number of trials, as it contains equal number of trials to “laser on” condition. Modulation along the vertical indicates a previous trial effect behavioral bias as a function of previous trial’s stimuli, for trials for which animals went Left, and were rewarded, therefore history of reward and choice is held fixed. Grey lines are different current trial [s_a_,s_b_] pairs, black line is average over pairs. **b**, similar to (a) for trials for which animals went Right and were rewarded. **c**, similar to (a) for all combinations of current and previous stimuli.

**Extended Data Fig. 9.**
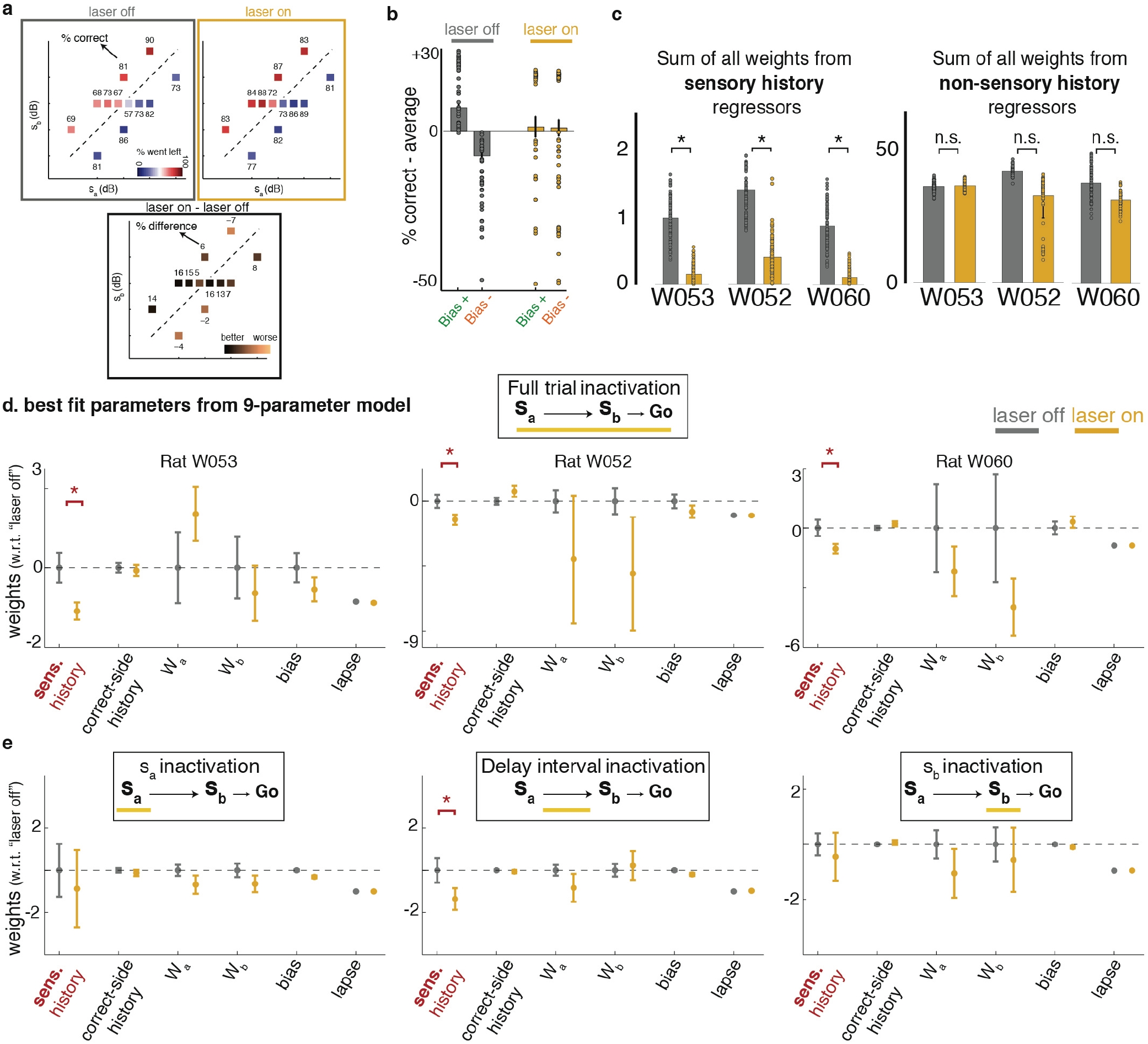
Optogenetics - Best fit parameters for non-sensory-history weights. **a**, Stimulus set and performance during optogenetic inhibition sessions, averaged over 44 sessions from 3 animals (delay interval of 2 second). Trials are grouped based on “laser off”, left, and “laser on”, right, conditions. The boxes represent the set of [s_a_, s_b_] pairs used in a session, with the color representing “% went left” and numbers above each box indicating “% correct”. The plot in the bottom shows the difference between “laser off” and “laser on” conditions, with positive values indicating improved performance on “laser on” condition and negative values indicating impaired performance. **b-c, similar** to Fig. 3d-f, with all data points overlaid on the bar-plots. b: n = 37 for each bar-plot (equal to the totall number of inactivation sessions). C: n = 600, from 200 iterations of 3-fold cross validation data; ‘*’ indicates p < 0.01 from one sided t-test. **d**, Best fit parameter values for all weights from the 9-parameter model (short-term **Sensory** history model, constrained version, figure 2d-e). Values are plotted as their mean once the average value from “laser-off”, condition is subtracted out. Except for the “sensory history”, none of the other weights were significantly affected by optogenetic inactivation of PPC. Errorbars show standard deviation of the mean. (n = 600, 200 iterations of 3fold CV; ‘*’ indicates p < 0.01 from one sided t-test). **e**, Similar to d, for period-selective optogenetic inhibitions, where PPC is selectively inhibited during first stimulus s_a_ (left), delay interval (middle), or second stimulus s_b_(right).

**Extended Data Fig. 10.**
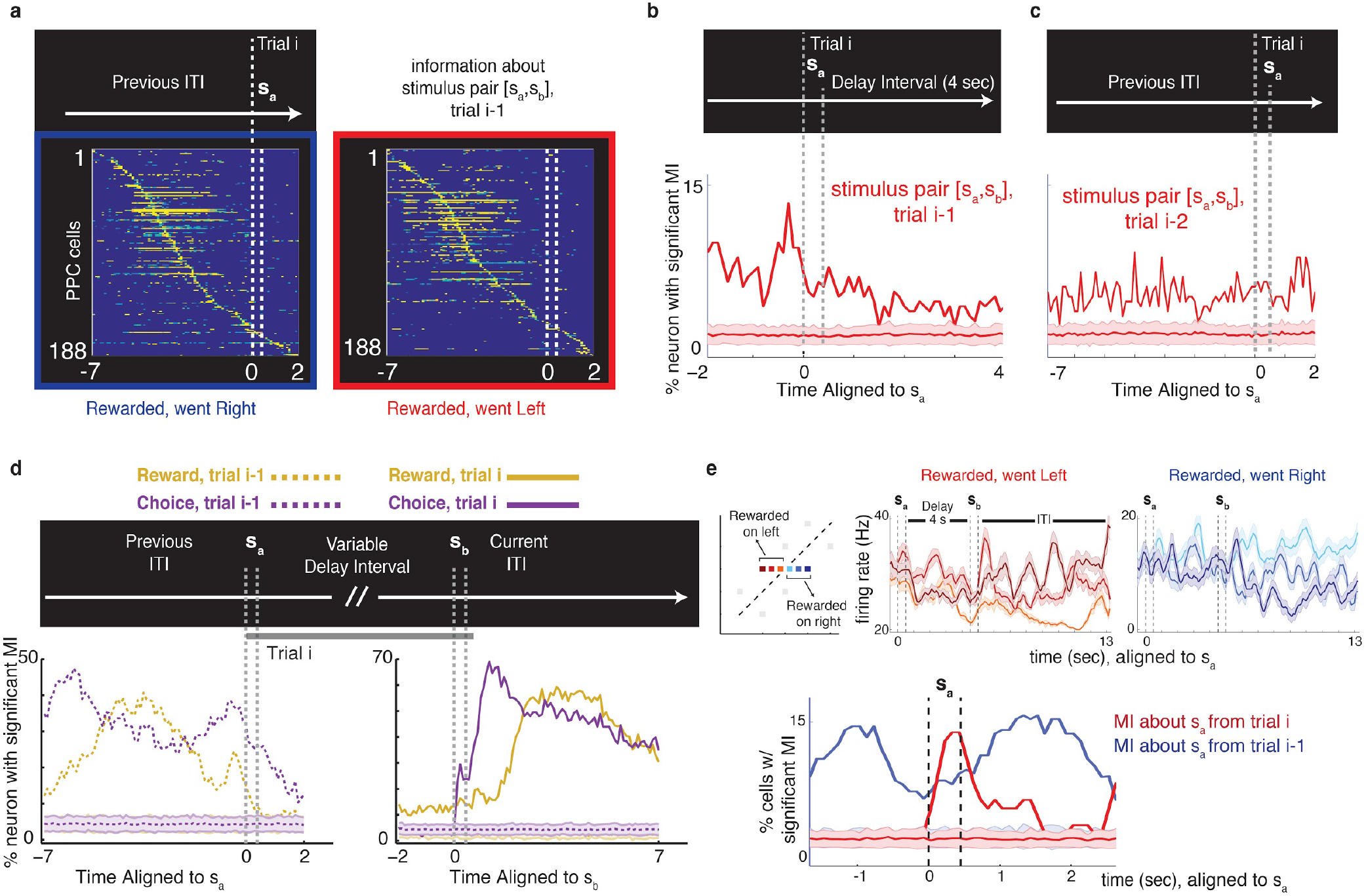
Mutual Information. **a, Sensory-history coding, 1 trial back, population analysis**, each row represents the time course of significant values of Mutual Information (MI) between cell’s firing rate and the stimulus pair [s_a_,s_b_] presented on the previous trial. Data from all trials with variable delay duration (minimum of 2 sec) was pooled and plots are aligned to the beginning of s_a_. Data from n=5 animals, and only cells with significant values of MI values are included. When estimating the MI, spurious information values can be caused by the inherent correlations between task parameters, like sensory stimuli and choice. To overcome this, conditional MI was calculated when only trials with same previous choice and reward status were considered, and sensory inputs were the only variable: Left panel, on the previous trial animals went right and were rewarded. Right panel, on the previous trial animals went left and were rewarded. **b, Sensory-history coding, 1 trial back**, % of cells with significant coding of stimuli presented on the previous trial (trial i-1), aligned to the start of trial i. Only trials with delay interval of larger 4 seconds are included in this analysis. **c**, **Sensory-history coding, two trials back**, % of cells with significant coding of stimuli presented two trials in the past (trial i-2), aligned to the start of trial i. **d**, % of cells with significant coding of animal’s choice and reward status, on both current trial (solid lines) and previous trial (dashed lines), when time is aligned to the current trial, either s_a_ (right), or s_b_ (left). **e**, In the standard stimulus set (Fig. 1b, [s_a_,s_b_] pairs along the diagonal lines), knowledge of the animal’s side choice, whether it was rewarded or not, and one of either s_a_ or s_b_ allows uniquely identifying the other stimulus (s_b_ or s_a_). Therefore, in order to probe whether neurons carried information for different values of s_a_ itself (as opposed to a combination, of choice, reward, and s_b_), we ran recording sessions with psychometric stimuli added to the standard stimulus set (top left panel). In this way, three different values of s_a_ are assigned to one fixed value of s_b_ and one fixed action (left in different shadows of red, and right in different shadows of blue). The firing rate of an example neuron is shown in response to different values of s_a_, only for trials in which animal responded left after Go cue (middle), or right (right), was rewarded, and the delay interval was 4 s. Even though choice, reward, and s_b_ are fixed, firing rates clearly differentiate values of s_a_. Bottom: summary of population analysis from psychometric recording sessions (as in the example in the upper panel), showing % cells with significant coding of s_a_ from trial i (red), or trial i−1 (blue, n = 142 cells).

